# The heritability of migration behaviours in a wide-ranging ungulate

**DOI:** 10.1101/2025.01.28.635302

**Authors:** Maegwin Bonar, Eric Wootton, Charles R. Anderson, George Wittemyer, Joseph M. Northrup, Aaron B. A. Shafer

**Affiliations:** Environmental & Life Sciences Graduate Program, Trent University, Peterborough, ON, K9L 0G2, Canada; Molecular Biology & Biochemistry Undergraduate Program, Trent University, Peterborough, ON, K9L 0G2, Canada; Mammals Research Section, Colorado Parks and Wildlife, Fort Collins, CO 80523, USA; Department of Fish, Wildlife and Conservation Biology, Colorado State University, Fort Collins, CO, 80523, USA; Wildlife Research and Monitoring Section, Ontario Ministry of Natural Resources, Peterborough, ON K9J 3C7, Canada

**Author notes:** Corresponding Author: Maegwin Bonar **Email:**. **Author Contributions:** MB, ABAS, JMN designed the study; JMN, CRA, GW secured funding for data collection and coordinated sample curation; MB, EW performed research and analyzed data; MB wrote the manuscript with input from ABAS, CRA, EW, GW, JMN. **Competing Interest Statement:** The authors declare no conflict of interest. **Classification:** Biological Sciences; Genetics.

**Keywords:** animal model, genomic relatedness matrix, heritability, migration

## Abstract

Migration behaviour is thought to be declining globally in the face of rapid human-mediated environmental change. While many species exhibit individual plasticity in their migratory behaviour, not all species demonstrate the level of plasticity necessary to adjust to novel conditions. Selection on heritable behaviours might therefore play an important role in the maintenance of migratory phenotypes for some species. Using GPS and genomic data from 242 individuals (256 animal-years) in a pedigree-free quantitative genetic approach, we estimated heritability, repeatability, and sources of environmental variation for migration traits in migrating mule deer (*Odocoileus hemionus*). We also estimated heritability of body size to ensure validity of our results. Heritability estimates of body size traits were comparable to the current estimates for ungulate body size. We found low heritability for broad patterns of migration timing, distance, and duration, but high heritability for movement rate along the migratory route. Our findings suggest that wild mule deer populations have the potential to respond to selection pressure generated by human activity or global environmental changes through microevolutionary changes in migration behaviours.

**Significance Statement:** Migration behaviour is critical for the reproduction and survival of a wide variety of taxa, yet there have been global declines in migrations in the face of rapid human-mediated environmental change. Despite our understanding that variation in migration behaviour has both genetic and environmental components, studies quantifying the sources of genetic variation contributing to migration phenotypes are lacking. Our study provides, to our knowledge, the first empirical evidence of heritability in a migration behaviour in ungulates. These results have implications for the evolution and maintenance of migration behaviours in natural populations.

## Introduction

Migration behaviour is critical for the reproduction and survival of a wide variety of taxa, yet populations have experienced global declines in migrations in the face of rapid human-mediated environmental change (1–4). While migratory behaviour is thought to be largely driven by environmental variation (5), behavioural variation is considered to have both a genetic component (6–8), and socially transmitted component (9). Migrating populations experiencing fluctuating environments can adapt to novel conditions through behavioural plasticity (10), and short-term evolution where selection acts on repeatable behavioural phenotypes that confer a fitness benefit (11). While many species exhibit individual plasticity in their migratory behaviour (10, 12), not all species demonstrate high levels of individual plasticity (13). Selection on heritable behavioural components might therefore play a stronger role in the maintenance of migratory phenotypes than previously thought. Meaning that populations would have the potential to respond to selection pressure generated by human activity or global environmental changes through microevolutionary changes in migration behaviours.

In this study, we used a pedigree-free quantitative genetic approach to quantify the proportion of phenotypic variance explained by additive genetic effects in spring migration behaviours in migrating mule deer (*Odocoileus hemionus*). In the spring, migration behaviour is linked to reproductive success as female mule deer migrate from their winter range to give birth on their summer range (14). Mule deer exhibit variation in components of their migration behaviour such as migration distance (13), and timing (15), and this variation could lead to variation in fitness. In the case of migration timing, mule deer typically migrate to match changes in resource availability and optimize migratory timing relative to both plant productivity and weather on their summer range (e.g. (16)). However, the fitness consequences of a mismatch in timing with spring green up may vary. For example, arriving ahead of spring green up and facing early spring storms could result in mortality, but in a milder year would mean first access to the best areas for giving birth. Similarly, migrating at a faster movement rate or greater distance could decrease exposure to predation along the route and lower mortality risk, but this would come at a cost of physical condition at an already taxing life-history stage.

Mule deer migrations are of interest to wildlife managers as recent anthropogenic development and climate change may threaten migratory routes (15, 16) and alter behaviours along the migratory route (17). While behavioural plasticity may be one way for at-risk populations to buffer against the effects of rapid environmental change, mule deer exhibit little to no migratory plasticity in terms of whether or where to migrate, meaning they could be less resilient to change (13). Consequently, adaptation to a rapidly changing environment might require microevolutionary change, for which a degree of heritability for migratory variation is required. We estimated heritability and sources of environmental variation (e.g., permanent environmental effects) in migration distance, timing, movement rate, and individual variation in weight, ingesta-free body-fat percentage, and body size measurements. Broadly, behaviour is often assumed to be highly plastic, suggesting that the magnitude of additive genetic variance should be low relative to the effect of environmental variation (18, 19).

Accordingly, we would expect the heritability of migration traits to be low compared to other traits such as morphological traits or life-history traits. A recent meta-analysis by Dochtermann et al. (20) reported the range of heritability of behavioural traits is 0.24–0.31. Although Dochtermann et al. (20) grouped migration and dispersal together in their models, they estimated moderate heritability of these traits (h^2^ = 0.46; 95% CI:0.33–0.57). Given our current understanding of the genetic and environmental drivers of migration (5), we predict a large proportion of phenotypic variation in migration behaviour to be explained by permanent environmental effects, and additive genetic effects (i.e., heritability) to explain a smaller proportion of phenotypic variance. Thus, we predict low heritability for migration timing, as variation in migration timing across ungulates is known to be heavily influenced by plant phenology (8) and environmental effects. We predict low heritability for migration distance, duration, and movement rate however heritability for these traits will likely be greater than that of migration timing as movements along the route are more constrained by physiological processes which have a genetic component. We predict high levels of repeatability for all migration traits given the large contribution of permanent environmental effects.

## Results

We captured 242 adult (>1 year old) female mule deer in the Piceance Basin of northwestern Colorado, USA (Figure 1). At the time of capture, individuals were fitted with store-on-board GPS collars, weighed, measured for chest girth and hind foot length, and blood samples were collected for genetic analysis. Of the 242 individuals captured, 207 had sufficient GPS data to determine a spring migration for at least one year from 2011-2016. We input GPS locations into Migration Mapper v3.0 (21) where we delineated migration routes visually for each individual using graphs of the individual’s net-squared displacement.

**Figure 1.**
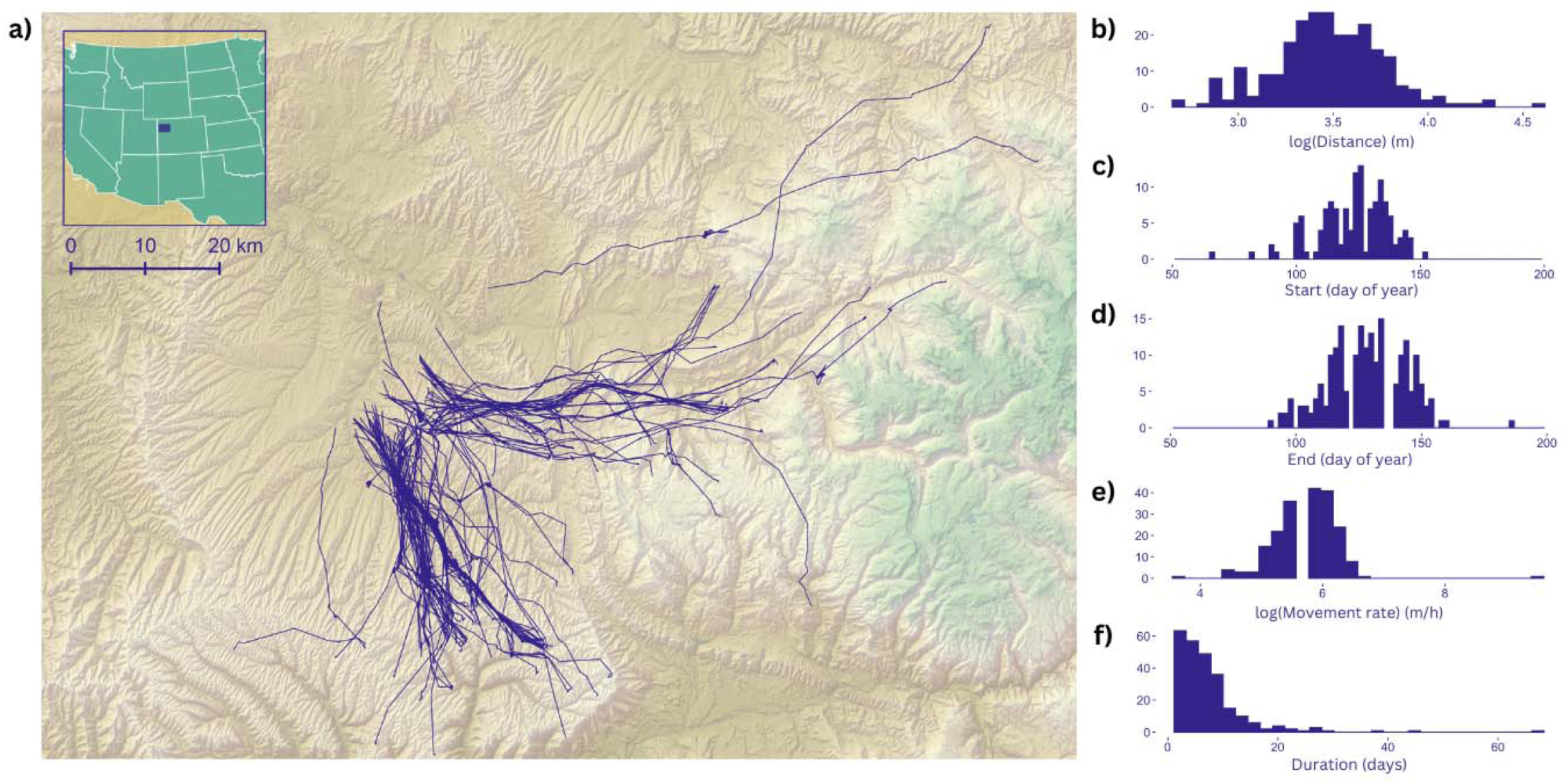
**a)** Mule deer migration routes for 256-animal years in the Piceance Basin, Colorado, USA. Frequency distributions of migratory traits for b) log-transformed Euclidian migration distance; c) migration start day; d) migration end day; e) log-transformed average movement rate along the route; and f) duration of migration.

The number of spring migrations per individual ranged from 1 – 4 (256 animal-years total). Frequency plots of migration traits are illustrated in Figure 1. The average start day of spring migration was May 2 (range March 8–June 1) and the average end day was May 10 (range April 1–July 6). The average duration of migration was 7 days (range 2–67). The average distance of migrations to the eastern summer range was 39.8km (range 14.2–94.9km) and the average distance to the southern summer range was 31.1km (range 14.7km–58.2km). The average movement rate for individuals was 327.1m/h (range 41.3–1,468.3m/h). Frequency plots of body size traits are available in the Supplementary Materials Figure S1. Body weight, hind leg length, chest girth, and ingesta-free body fat percentage does not differ significantly between east-west and north-south migrating individuals (22) and there is no genetic subdivision within the population (15, 22).

After applying quality filtering, 143 individuals were retained in the unfiltered genomic relatedness matrix (GRM). The genomic relatedness matrix included 10,097 SNPs and 20,306 pairwise relatedness coefficients, of which 104 pairs had a relatedness coefficient higher than 0.1 (Figure S3, Table S1). The results presented herein for the models that used the GRM with a relatedness cutoff of 0.1; this GRM included 109 individuals, had a mean diagonal of 1.02, and the total variance of the relatedness coefficient of the off-diagonal was 0.0001. We also performed all analyses with an unfiltered GRM, and a GRM with a relatedness cutoff of 0.05. The results of those models can be found in the Supplementary Materials.

Narrow-sense heritability h^2^ (CI) of body size traits ranged from 0.12 (0.01-0.57) for ingesta-free body-fat (measurements taken in March of the year of migration) to 0.27 (0.06-0.53) for hind leg length. Permanent environmental effects pe^2^ (CI) and repeatability (i.e., among-individual differences, ind^2^ (CI)) were similarly low for ingesta-free body-fat 0.07 (0.01-0.31) and 0.19 (0.03-0.66). Permanent environmental effects for other body size traits ranged from 0.22 (0.05-0.63) for weight to 0.45 (0.17-0.84) for hind leg length. Repeatability estimates were all moderate for the remaining body size traits with 0.42 (0.14-0.86) for weight, 0.56 (0.27-0.88) for chest girth, and 0.72 (0.46-0.93) for hind leg length (Figure 1, Table 2).

Heritability for start day and end day of migration was low, with values of 0.04 (0.01-0.11) for start day, and 0.05 (0.01-0.14) for end day. Variation in migration start and end day was explained primarily by permanent environmental effects which were captured in the year-to-year variation (Figure 1, Table 2).

Movement behaviours along the migration route ranged considerably, with distance and duration having lower heritability (0.14 (<0.01-0.66) and 0.10 (<0.01-0.37) respectively). Average movement rate was highly heritable with a value of 0.76 (0.12-0.94). Variation in distance and duration was explained largely by individual identity (Figure 2).

**Figure 2.**
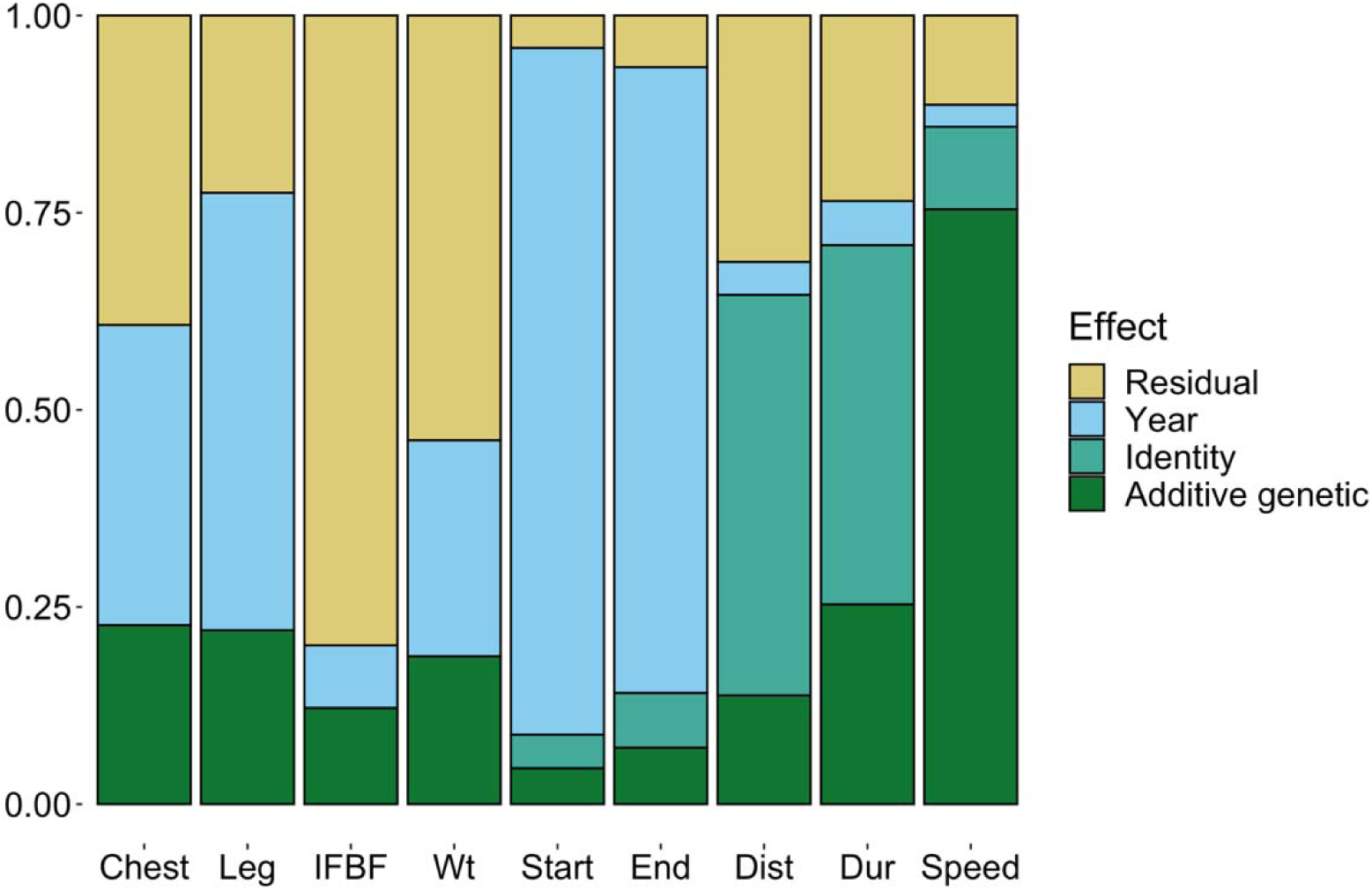
Variance partitioning of body size and migration behavioural traits. Proportion of phenotypic variance of traits explained by additive genetic effects, permanent environmental effects (year and identity), and residual effects.

## Discussion

Migratory behaviour is a complex process that is thought to emerge from a combination of physiological, morphological, and cognitive traits (5, 8), suggesting that genetics at least partially underpin the phenotypic variation of migratory traits. Knowing the general pattern of heritability of migration behaviour is important as inference on the selection, evolution and maintenance of migration behaviour often assumes that phenotypic differences among groups (e.g., age, sex, social environment, populations, species) are adaptive and driven by underlying genetic differences (i.e., the phenotypic gambit; (23)).

However, whether this is true, or not, is unclear for many taxa yet critically important as the degree of heritability of different traits could provide insight to which traits are most plastic under ongoing global environmental change. Our study provides, to our knowledge, the first empirical evidence of heritability in migration behaviour in ungulates. We found low heritability for broad patterns of migration timing, distance, and duration, but high heritability for movement rate along the migratory route. Additionally, our heritability estimates of body size traits were comparable to the current estimates of ungulate body size (e.g. (24–26)). Therefore, we have some confidence that our pedigree-free approach to estimating heritability of both body size and migration behaviours has likely captured a true signal. These findings have important implications for our understanding of the evolution of migration in ungulates and for how migration behaviours may be influenced by human-mediated environmental change.

### High heritability of movement rate, low heritability of migration timing

Contrary to our prediction, we found high heritability for average movement rate during migration (h^2^ = 0.76) after accounting for age, body size and distance migrated. Heritability of migration behaviour in ungulates has not been measured in other systems, however Gervais et al. (27) quantified heritability of movement rate and space-use behaviours within home ranges of roe deer (*Capreolus capreolus*) and found a moderate heritability (h^2^ = 0.21 ± 0.08) for average daily movement speed and high heritability of distance to roads (h^2^ = 0.70 ± 0.11). The genetic basis for speed and stamina has been well documented in racehorses, with genome-wide scans identifying candidate loci correlating with speed (28, 29).

Heritability estimates of speed in racehorses varies within the literature from low (h^2^ range: 0.074 ± 0.012 – 0.124 ± 0.006; (30)) to moderate and high (h^2^ range: 0.38 ± 0.03 – 0.68 ± 0.05; (31)). A genetic basis for movement traits in mule deer has been suggested by Lingle (32) who showed that the escape gaits of mule deer, white-tailed deer, and their F1 hybrid crosses were highly consistent. High heritability does not necessarily imply a strong response to selection, due to the lack of independence between additive genetic variance and environmental effects (33). Social environment and maternal genetic effects may play a large role in shaping movement rate as mule deer typically follow their mother’s migration route (15, 34). In the case of movement rate, physiological constraints, and reduced selection pressure on movement rate as a trait are likely to limit response to selection.

Low heritability can result from the erosion of additive genetic variance by stabilizing or directional selection (35), or from increased residual variance due to environment (36). Variation due to year explained the largest proportion of phenotypic variation for migration start and end day in our models.

Work in this system by Lendrum et al. (16) found that spring migration in mule deer differed among years and was related to the environmental variables of snow depth, and plant phenology. Similarly, Monteith et al. (37) found the same patterns of year-to-year variation in migration timing, linked to southern oscillation index, snow depth and plant phenology for mule deer in the Sierra Nevada. Migration timing across ungulates is known to be influenced by plant phenology (e.g., elk (*Cervus elaphus canadensis*) (38), mule deer (16, 37) red deer (*Cervus elaphus*) (39), roe deer (39, 40)), with species exhibiting plasticity in their migration timings in order to optimize foraging along their route. Plant phenology is tightly linked to changes in temperature (41) which contribute to yearly variation in spring green-up. Low heritability of migration timing is in line with our prediction that most of the phenotypic variation in migration timing is explained by permanent environmental effects. That being said, the high heritability of movement rate suggests that the rate at which mule deer migrate has the potential change to some degree at the population level. Under rapid environmental change it may be adaptive for migrations to become faster when conditions become less predictable.

### Individual repeatability of migration distance and duration

All heritability values estimated were substantially lower than our estimates of individual repeatability, which is assumed to set the upper bound for heritability (42, 43). High estimates of repeatability for migration behaviours fit with our prediction, given the large contribution of permanent environmental effects to phenotypic variance and the fidelity that mule deer in this system show to their summer and winter ranges. Bell et al. (42) found in their meta-analysis the average measure of repeatability of behaviour was 0.37. Average repeatability across migration traits was 0.67, and individual identity largely explained variation in migration distance and duration. High repeatability of migration behaviours might be linked to strong spatial fidelity exhibited by mule deer (16, 44, 45). For example, Sawyer et al. (13) found high levels (>80%) of site fidelity in mule deer migration routes, regardless of age, reproductive status, or number of years monitored and Northrup et al. (45), working in our system, showed high year-to-year fidelity in summer and winter ranges indicating they migrate to and from the same locations annually.

Similarly, Mahoney and Schaefer (46) found that rank order of caribou (*Rangifer tarandus*) migration was highly consistent among individuals, and also independent of sex or age. Studies reporting consistent among-individual differences in movement and space-use behaviours are becoming more prevalent across taxa (27, 47, 48), however the mechanisms underpinning individual repeatability of behaviours is still in need of exploration. Sawyer et al. (39) proposed that the high level of consistency in mule deer migration behaviour is due to the strong reliance on memory for navigation. Migrations shaped largely by memory and experience may be less flexible than those acquired by social learning from conspecifics (49, 50).

### Considerations for free-ranging populations

When estimating heritability using GRM-based approaches, sample size and variance in relatedness are important factors. Small sample size can increase sampling variance, which would bias V_A_ downwards, thereby decreasing heritability estimates (51). Our study and that of Gervais et al. (27) are two examples which demonstrate a GRM-based approach can be used to estimate heritability using small samples sizes of a few hundred unrelated individuals. The GRM variance of relatedness for our study was low, because like other studies of wild ungulates (e.g., (27, 52) our GRM was skewed towards unrelated individuals. With SNP data, the GRM variance is inversely proportional to the variance of h^2^ (53), however, by removing highly related individuals, we can reduce error resulting from being unable to sample the full spectrum of relatives (54). However, population-wide estimates using unrelated individuals can lead to an underestimation of heritability estimates compared to pedigree approaches (55). While these factors could bias our estimates of heritability low, we can be more confident that our higher heritability estimates are capturing truly heritable traits.

GRM-based approaches make estimating heritability of wild populations more accessible, however one of the challenges it presents is understanding the effect of heterogeneous landscapes on heritability estimates. Observed heritability of a trait may be partially due to related individuals sharing more similar habitats, which would result in the genetic and environmental sources of variation being potentially confounded (56). In our system relatives are more likely to share similar environments as offspring are philopatric to their mother. Environmental similarity was shown to affect heritability estimates in red deer populations (57) but did not have substantial effect in Soay sheep (*Ovis aries*; (58)). However, if closely related individuals are excluded from the analysis, which is the case in our study, this source of bias should be reduced because among distantly related individuals, genomic similarity is poorly correlated with pedigree relationship, and it is only the pedigree relationship that might be correlated with environmental similarity (55).

Our study provides empirical demonstration of heritability of a migration trait. Broad patterns of migration timing have low heritability and are heavily influenced by year-to-year environmental variation, while movement behaviour during migration is more heritable and subject to individual differences in repeatability. Future studies should seek to quantify the heritability of more movement and space-use behaviours that affect fitness along the migration route, such as tracking forage, and habitat selection or avoidance of anthropogenic features. Changes to the landscape resulting from human activity are having a negative effect on migration routes globally, often resulting in the disappearance of migration routes (1, 2). Bonnet et al. (59) found in their recent study that across bird and mammal populations, estimates of additive genetic variance of fitness are often substantial and more than expected; and suggests that the adaptive evolution may occur on a generation-to-generation time scale. Our findings imply that wild mule deer populations have the potential to respond to selection pressure generated by human activity or global environmental changes through microevolutionary changes in migration behaviours.

## Materials and Methods

Information on the study site can be found in the Supplementary Methods.

### Individual sampling and phenotypes

Adult (>1 year old) female mule deer were captured using helicopter net gunning and fitted with store-on-board GPS radio collars (Advanced Telemetry Systems, Isanti MN, USA) with three different relocation schedules (5 hours, 60 minutes, and 30 minutes) depending on the individual. See Supplementary Methods for details on deer capture and processing. All procedures were approved by the Colorado State University (protocol ID: 10-2350A) and Colorado Parks and Wildlife (protocol ID: 15-2008) Animal Care and Use Committees.

GPS locations were input into Migration Mapper v3.0 (21) where we removed erroneous locations, and then delineated migration routes visually for each individual using graphs of the individual’s net-squared displacement. Some individuals were collared for more than one year, so we determined spring migrations for every year in which data were available. After migration routes had been determined, we recorded the start and end of spring migration in day of year, the duration of the migration in days, and calculated Euclidean distance from the first GPS location of the migration route to the last GPS point. Movement rate was calculated per single step length, and then averaged across all step lengths for the entire route. To account for the three different relocation schedules, we aggregated relocations for all individuals to 5-hour intervals before calculating movement rate. Habitat characteristics (i.e., vegetation type) occurring along the migratory paths of mule deer was similar across the study areas (60), however individuals that migrated east-west on average travelled greater distances than individuals migrating north-south.

### DNA extraction and library preparation for RAD sequencing

We extracted DNA from blood samples using the DNeasy Blood and Tissue Kit (Qiagen, Inc., Valencia, CA, USA), following the manufacturer’s protocol. We generated restriction site-associated DNA sequencing (RADseq) libraries using an adapted protocol from Parchman et al. (61) and Peterson et al. (62). See Supplementary Methods for details on lab protocols. The libraries were sequenced at The Centre for Applied Genomics (TCAG) in The Hospital for Sick Children (SickKids) on an Illumina HiSeq 2500 to produce 2 × 126 base pair paired-end reads.

### Bioinformatic pipeline for RADseq data

Full details on packages used in the bioinformatic pipeline can be found in Supplementary Materials. Briefly, Fastq files were demultiplexed within the Stacks v2.3 module (63), and aligned against the white-tailed deer genome (*Odocoileus virginianus*) (Accession No. JAAVWD000000000) that was recently annotated (64). Mapped reads were sorted and indexed using SAMtools (65), and then we produced a variant call format (VCF) file using Stacks. We also only retained one single nucleotide polymorphism (SNP) per locus to meet the assumptions of linkage equilibrium in subsequent analyses. The VCF file was converted into a binary fileset using PLINK v.19 (66) and that was used to generate the classic genomic relatedness matrix (GRM) in GCTA v1.92.4 (67). Highly related individuals were removed at two levels (r > 0.10, and r > 0.05) to reduce heterogeneity in the matrix. All individuals were retained to generate the unfiltered GRM even if phenotypic data were missing as this improves relatedness estimates. All subsequent analyses were run the unfiltered and filtered GRMs respectively.

### Quantitative analyses

We used the animal model approach to partition the phenotypic variance of migration and body size traits (V_P_) into additive genetic variance (V_A_), permanent environmental effects (V_PE_) and residual variance (V_R_) such that V_P_ = V_A_ + V_PE_ + V_R_ conditional on any fixed effects. We included age and month of capture (December or March) as fixed effects in all body size models as body size may differ between life-history stages and time of year. There was little variation in pregnancy rates as 95% of the deer captured in March were pregnant at the time. Table 1 summarizes the fixed and random effects for each animal model. Briefly, we included age and log-transformed hind leg length in all the migration models; summer range was included as a fixed effect for start day, end day, distance, and duration models; log-transformed distance of migration route was included in the movement rate and duration models. We included year as a random effect in all the models to account for interannual differences in individual behaviour and environmental conditions. Individual identity was included as a random effect in all the migration models to account for multiple migrations per individual. We included the GRMs in the models as a random effect as our measure of pairwise relatedness. The unfiltered GRM was nonpositive definite but could nevertheless be implemented in an animal model by bending the matrix (68).

**Table 1.** Summary of fixed and random effect for each animal model. All in the animal models require a measure of pairwise relatedness as a random effect, our measure of relatedness is determined using genomic relatedness matrices (GRMs). We log-transformed response variables with continuous distributions to meet assumptions of homoscedasticity and normality of residuals and used a Poisson distribution for the discrete response variables.

| Model | Response variable | Fixed effects | Random effects | Model distribution |
| --- | --- | --- | --- | --- |
| 1 | Chest girth | age + month of capture | relatedness (GRM) + year | Normal |
| 2 | Hind leg length | age + month of capture | relatedness (GRM) + year | Normal |
| 3 | Ingest-free body-fat | age + month of capture | relatedness (GRM) + year | Normal |
| 4 | Weight | age + month of capture | relatedness (GRM) + year | Normal |
| 5 | Start day | age + summer range + log(hind leg length) | relatedness (GRM) + year + identity | Poisson |
| 6 | End day | age + summer range + log(hind leg length) | relatedness (GRM) + year + identity | Poisson |
| 7 | Distance | age + summer range + log(hind leg length) | relatedness (GRM) + year + identity | Normal |
| 8 | Duration | age + summer range + log(hind leg length) + log(distance) | relatedness (GRM) + year + identity | Poisson |
| 9 | Movement rate | age + log(hind leg length) + log(distance) | relatedness (GRM) + year + identity | Normal |

**Table 2.** Variance component estimates and their associated ratios for body size and migration traits.

| Trait | $^1h^2$ | $pe^2$ | $ind_2$ | $^1V_A$ | $V_{ID}$ | $V_{YR}$ | $V_R$ |
| --- | --- | --- | --- | --- | --- | --- | --- |
| Chest girth | 0.25 (0.06-0.58) | 0.30 (0.09-0.72) | 0.56 (0.27-0.88) | 0.001 (<0.001-0.002) | - | 0.001 (<0.001-0.006) | 0.001 (0.001-0.002) |
| Hind leg length | 0.27 (0.06-0.53) | 0.45 (0.17-0.84) | 0.72 (0.46-0.93) | <0.001 (<0.001-0.001) | - | 0.001 (<0.001-0.005) | <0.001 (<0.001-0.001) |
| Ingest-free body-fat | 0.12 (0.01-0.57) | 0.07 (0.01-0.31) | 0.19 (0.03-0.66) | 0.009 (0.001-0.044) | - | 0.006 (<0.001-0.030) | 0.060 (0.027-0.085) |
| Weight | 0.20 (0.03-0.63) | 0.22 (0.05-0.63) | 0.42 (0.14-0.86) | 0.002 (<0.001-0.007) | - | 0.003 (<0.001-0.015) | 0.007 (0.002-0.0100) |
| Start day | <sup>a</sup>0.08 (0.01-0.23)<br><sup>b</sup>0.04(0.01-0.11) | 0.86 (0.62-0.98) | 0.93 (0.78-0.99) | <sup>a</sup>0.001 (<0.001-0.002) <sup>b</sup>13.37 (3.82-33.96) | 0.001 (<0.001-0.002) | 0.017 (0.003-0.069) | 0.001 (<0.001-0.002) |
| End day | <sup>a</sup>0.11 (0.01-0.32) <sup>b</sup>0.05 (0.01-0.14) | 0.79 (0.51-0.97) | 0.90 (0.71-0.99) | <sup>a</sup>0.001 (<0.001-0.002) <sup>b</sup>16.43 (4.49-41.56) | 0.001 (<0.001-0.002) | 0.011 (0.002-0.045) | 0.001 (<0.001-0.002) |
| Distance | 0.14 (<0.01-0.66) | 0.55 (0.04-0.88) | 0.68 (0.17-0.92) | 0.017 (0.001-0.087) | 0.063 (0.002-0.123) | 0.005 (<0.001-0.027) | 0.039 (0.011-0.104) |
| Duration | <sup>a</sup>0.25 (<0.01-0.94) <sup>b</sup>0.10 (<0.01-0.37) | 0.51 (0.02-0.98) | 0.76 (0.07-1.00) | <sup>a</sup>0.084 (0.001-0.347) <sup>b</sup>4.80 (0.04-20.86) | 0.151 (0.001-3.88) | 0.019 (0.001-0.113) | 0.078 (0.001-0.321) |
| Movement rate | 0.76 (0.12-0.94) | 0.13 (0.01-0.72) | 0.88 (0.65-0.97) | 0.293 (0.047-0.448) | 0.041 (0.001-0.274) | 0.011 (0.001-0.059) | 0.044 (0.012-0.134) |

<sup>1</sup>For Poisson distributed traits (start day, end day, duration) heritability and additive genetic effect values are given in both the latent model state <sup>a</sup> and true data state <sup>b</sup>.

We fitted the animal models using a Markov Chain Monte Carlo for generalized linear mixed models with the MCMCglmm package in R (69). We ran the algorithm 4 times for each model, resulting in 4 chains, and thinned the chains at an interval of 100. We used a burn-in period of 10,000 iterations and a total of 500,000 iterations per chain to estimate the posterior distribution. We decided to log-transform traits with continuous distributions to ensure proper support. We ran the start day, end day, and duration models using a Poisson distribution. Narrow sense heritability of the traits was estimated as h^2^ = V_A_/(V_A_ + V_PE_ + V_R_) and the permanent environmental effect as pe^2^ = V_PE_/(V_A_ + V_PE_ + V_R_). We measured individual repeatability as the ratio of among-individual variance (genetic and nongenetic) over the total phenotypic variance ind^2^ = V_A_+V_PE_/ (V_A_ + V_PE_ + V_R_) (70). For the models using the Poisson distribution we report the heritability and additive genetic variance on both the latent scale and true data scale which were determined using the R package QCglmm (71).

## Acknowledgments

We respectfully acknowledge the territory where data were collected as the ancestral homelands of the Ute peoples. Data analyses were conducted at Trent University, which is on the traditional territory of the Mississauga Anishinaabeg. This work was supported by Natural Sciences and Engineering Research Council of Canada (NSERC) Discovery Grant (ABAS and JMN); NSERC Vanier PhD Fellowship (MB); Compute Canada Resources for Research Groups (ABAS). Mule deer capture and monitoring was funded by Colorado Parks and Wildlife (CPW), ExxonMobil Production/XTO Energy, Williams Companies/WPX Energy, Shell Exploration and Production, EnCana Corp., Marathon Oil Corp., Federal Aid in Wildlife Restoration (W-185-R), the Colorado Mule Deer Foundation, the Colorado Mule Deer Association, Safari Club International, Colorado Oil and Gas Conservation Commission, and the Colorado State Severance Tax. We thank D. Freddy for developing the field research effort and L. Wolfe, M. Fisher, C. Bishop, D. Finley (CPW) and numerous field technicians for capture expertise and field assistance. We thank Quicksilver Air, Inc. for deer captures, and L. Gepfert (CPW) and Coulter Aviation, Inc. for fixed-wing aircraft support.

## Notes

### Competing Interest Statement

The authors have declared no competing interest.

