## Supplemental Appendix for "The heritability of migration behaviours in a wide-ranging ungulate"

**Supplementary Materials**

**Supplemental Methods:**

**Study site**

The climate in the Piceance Basin of northwestern Colorado, USA was characterized by cold winters (range = −37.2–22.8°C) and warm dry summers (range: −2.2–35.6°C) with monsoonal precipitation in the late summer (1). The area was topographically variable with the dominant vegetation consisting of big sagebrush (*Artemisia tridentata*) and a pinyon pine (*Pinus edulis*)–Utah juniper (*Juniperus osteosperma*) shrubland complex. Other dominant shrubs included Utah serviceberry (*Amalenchier utahensis*), mountain mahogany (*Cercocarpus montanus*), bitterbrush (*Purshia tridentata*), and mountain snowberry (*Symphoricarpos oreophilus*). This area was popular for hunting during the fall with an annual average of 511 deer harvested in the wildlife management unit (Game Management Unit 22), which encompassed the entire study area (1). Mule deer in this area typically occupy their winter range between October and April of each year (2, 3) and migrate to two different summer ranges (2) located south or east of the winter range (Figure 1). Elevation on the winter study area ranged from 1,675 to 2,285 m and from 2,000 to 2,800 m on the summer study areas.

**Individual capture and sampling**

Deer were spotted visually by the helicopter capture crew and captured using a net gun. Deer were then blindfolded, hobbled, and administered 0.5 mg/kg of Midazolam and 0.25 mg/kg of Azaperone intramuscularly to alleviate capture-related stress (dose of both drugs based on an average weight of 75 kg). Deer were transported to a central processing site typically within 2 km of the capture site, where they were weighed, measured for chest girth and hind foot length, and blood samples were collected for genetic analysis. We estimated their age using tooth replacement and wear (4–6). We also obtained a body condition score by palpating the rump and measured the thickness of subcutaneous rump fat and the depth of the longissimus dorsi muscle using ultrasound (7–11). We used the body condition score and ultrasound measurements to estimate the percent ingesta‐free body fat of each deer in March of the year of capture (10, 11). During late‐winter captures, we assessed pregnancy using ultrasound and for does for which we did not detect a fetus, we confirmed pregnancy status using pregnancy‐specific protein B from blood samples. Deer were released at the processing site immediately following blood sample collection and GPS collar attachment.

**Library preparation for RAD sequencing**

Samples were incubated, digested overnight, and heat-inactivated in 96-well plates (see Haworth et al (12) for reaction conditions and primer details). Each 96-well plate had UltraPure distilled water (Invitrogen, 1897011) as negative controls. Restriction digested DNA was combined with 7 μl of ligation mixture and 3 μl of one of the 24 available Sbfl adapters (1.0 μM), and adapters were ligated at 16°C for 3 hours. We purified DNA fragments of artifacts following manufacturer protocol for AMPure XP beads (Beckman Coulter, A63880). Adapter-ligated fragments were amplified in four separate 10 μl reactions that incorporated barcodes. We pooled and purified samples following manufacturer protocol for QIAquick PCR Purification kit (Qiagen, 28106) for a final elution to 42 μl. We performed size selection between 450 bp to 700 bp on 80 μl purified libraries and performed gel purification following manufacturer protocol for QIAquick Gel Extraction kit (Qiagen, 28706) for a final elution to 60 μl. We characterized the pooled and purified final libraries with on a TapeStation using the D1000 kit (Agilent, 5067-5582).

**Bioinformatic pipeline for RADseq data**

Fastq files were demultiplexed using *process_radtags* within the Stacks v2.3 module (13). Parameters within *process_radtags* included the removal of any read with an uncalled base and the discarding of reads with low-quality scores. The demultiplexed sample files were aligned against the white-tailed deer genome. Mule deer and white-tailed deer can hybridize (14) so we opted to use the long-read-based draft genome of white-tailed deer (*Odocoileus virginianus*) (Accession No. JAAVWD000000000) that was recently annotated (15); we note that the mule deer genome available at the time of analysis was simply a consensus sequence from reads mapped to earlier versions of the white-tailed reference (16). Mapped reads were sorted and indexed using SAMtools (17). We then ran the *gstacks* and *populations* program within STACKs, retaining loci found in at least 80% of samples (r = 0.80), with a minor allele frequency of 1% (min_maf = 0.01), and heterozygosity upper bound of 0.8 (max_het = 0.8) that produced a variant call format (VCF) file. We also only retained one single nucleotide polymorphism (SNP) per locus (--write_single_snp), to meet the assumptions of linkage equilibrium in subsequent analyses.

The VCF file was filtered using PLINK v.19 (18) to only include individuals with less than 10% missing data. Using VCFtools (19) we determined F_IS_ and observed and estimated homozygosity, and removed outlier individuals with F_IS_ < -0.1 on the basis of excess heterozygosity (20, 21), which was visualized with a principal component analysis (Figure S1, S2) using PLINK and R v.4.2.1 (22). The VCF file was converted into a binary fileset using PLINK and that was used to generate the classic genomic relatedness matrix (GRM) in GCTA v1.92.4 (23). We examined the diagonal values of the GRM to identify potential problematic samples with inflated self-relatedness values and used this to further inform filtering. Highly related individuals were removed at two levels (r > 0.10, and r > 0.05) to reduce heterogeneity in the matrix using the grm-cutoff flag. All individuals were retained to generate the unfiltered GRM even if phenotypic data were missing as this improves relatedness estimates. All subsequent analyses were run the unfiltered and filtered GRMs respectively.

**Supplementary Figures and Tables:**

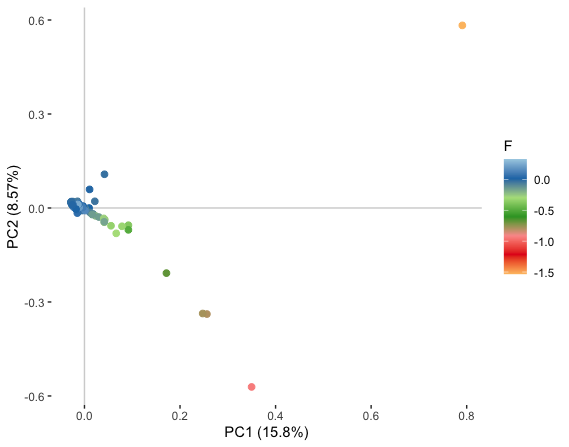

**Figure S1.** Principal coordinate analysis of individuals by F_IS_ coefficient, prior to removal based on excess heterozygosity (n = 155).

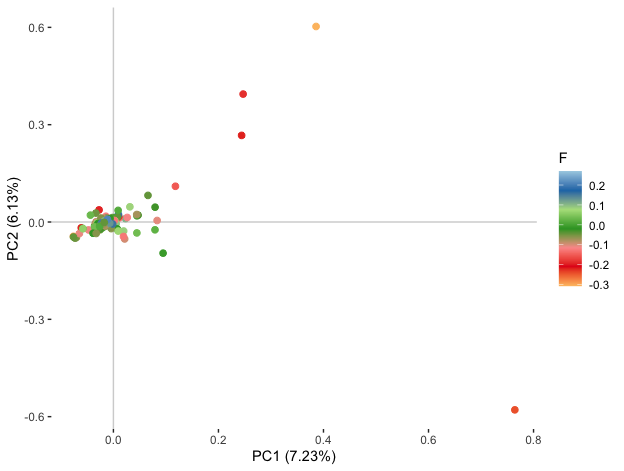

**Figure S2.** Principal coordinate analysis of individuals by F_IS_ coefficient, after removal based on excess heterozygosity (n = 143).

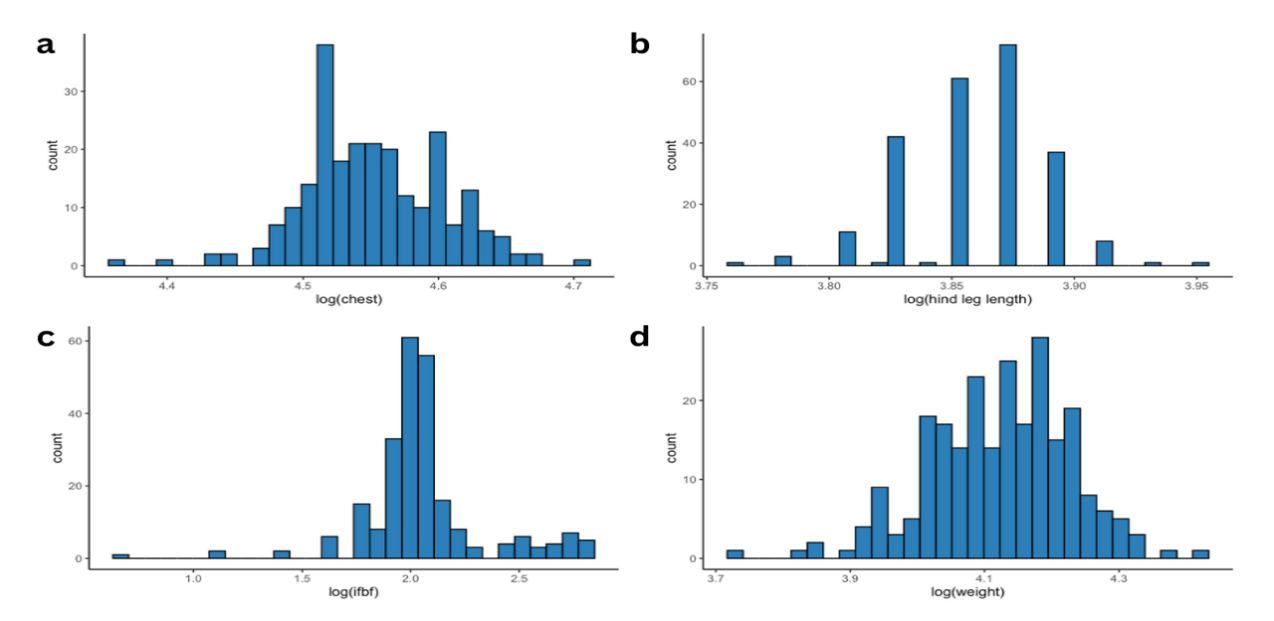

**Figure S3.** Frequency distributions of body size traits for 242 individuals. Body size traits are a) chest girth (cm); b) hind leg length (cm); c) ingesta-free body fat (%); and d) weight (kg).

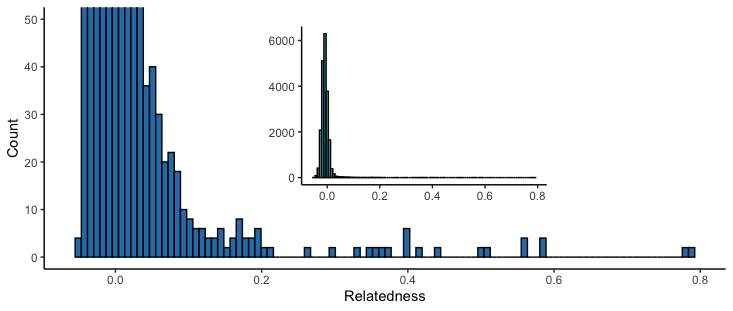

**Figure S4**. Distribution of pairwise genomic relatedness coefficients (n = 20,306).

**Table S1.** Summary data for unfiltered and filtered genomic relatedness matrices (GRM). The GRM was filtered based on two relatedness coefficient cut-offs, 0.1 and 0.05.

| **GRM** | **N** | **SNPs** | **mean off-diagonal** | **var off-diagonal** | **mean diagonal** | **var diagonal** |
| --- | --- | --- | --- | --- | --- | --- |
| unfiltered | 143 | 10,097 | -0.0071543 | 0.00072768 | 0.9904386 | 0.06703364 |
| 0.1 cutoff | 109 | 10,097 | -0.0074564 | 0.0001735 | 1.0177661 | 0.08338047 |
| 0.05 cutoff | 88 | 10,097 | -0.0072232 | 0.00012267 | 0.9916709 | 0.05397322 |

**Table S2.** Variance component estimates and their associated ratios for body size and migration traits using the unfiltered GRM. h^2^: heritability, pe^2^: permanent environmental effects, ind^2^: repeatability, V_A_: additive genetic variance, V_ID_: variance due individual identity, V_YR_: variance due to year, V_R_: residual variance.

| **Trait** | **^1^h^2^** | **pe^2^** | **ind^2^** | **^1^V_A_** | **V_ID_** | **V_YR_** | **V_R_** |
| --- | --- | --- | --- | --- | --- | --- | --- |
| Chest girth | 0.28 (0.07-0.60) | 0.32 (0.10-0.73) | 0.60 (0.30-0.89) | 0.001 (<0.001-0.002) |  | 0.001 (<0.001-0.006) | 0.001 (<0.001-0.002) |
| Hind leg length | 0.27 (0.07-0.51) | 0.47 (0.19-0.85) | 0.74 (0.50-0.94) | <0.001 (<0.001-0.001) |  | 0.001 (<0.001-0.005) | <0.001 (<0.001-0.001) |
| Ingest-free body-fat | 0.13 (0.01-0.60) | 0.06 (0.01-0.26) | 0.19 (0.03-0.67) | 0.008 (0.001-0.039) |  | 0.004 (<0.001-0.020) | 0.049 (0.022-0.067) |
| Weight | 0.23 (0.03-0.65) | 0.20 (0.04-0.60) | 0.43 (0.11-0.85) | 0.002 (<0.001-0.007) |  | 0.003 (<0.001-0.012) | 0.006 (0.002-0.009) |
| Start day | ^a^0.06 (0.01-0.20) ^b^0.03 (0.01-0.09) | 0.88 (0.68-0.98) | 0.94 (0.82-0.99) | ^a^0.001 (<0.001-0.002) ^b^11.16 (3.5-26.87) | 0.001 (<0.001-0.073) | 0.018 (0.003-0.073) | 0.001 (<0.001-0.002) |
| End day | ^a^0.09 (0.01-0.26) ^b^0.04 (0.01-0.11) | 0.83 (0.59-0.98) | 0.92 (0.77-0.99) | ^a^0.001 (<0.001-0.002) ^b^13.63 (4.16-32.69) | 0.001 (<0.001-0.002) | 0.012 (0.002-0.051) | 0.001 (<0.001-0.002) |
| Distance | 0.13 (0.01-0.56) | 0.56 (0.06-0.87) | 0.70 (0.24-0.91) | 0.014 (0.001-0.062) | 0.056 (0.003-0.099) | 0.004 (<0.001-0.022) | 0.032 (0.010-0.081) |
| Duration | ^a^0.17 (<0.01-0.83) ^b^0.07 (<0.01-0.34) | 0.49 (0.02-0.98) | 0.66 (0.05-1.00) | ^a^0.049 (0.001-0.249) ^b^2.5 (0.04-13.4) | 0.123 (0.001-0.308) | 0.012 (0.001-0.071) | 0.093 (0.001-0.288) |
| Movement rate | 0.75 (0.37-0.92) | 0.10 (0.01-0.45) | 0.86 (0.64-0.95) | 0.243 (0.117-0.350) | 0.024 (0.001-0.133) | 0.001 (<0.001-0.051) | 0.045 (0.016-0.111) |

^1^For Poisson distributed traits (start day, end day, duration) heritability and additive genetic effect values are given in both the latent model state ^a^ and true data state ^b^

**Table S3.** Variance component estimates and their associated ratios for body size and migration traits using the GRM with 0.05 relatedness cut-off. h^2^: heritability, pe^2^: permanent environmental effects, ind^2^: repeatability, V_A_: additive genetic variance, V_ID_: variance due individual identity, V_YR_: variance due to year, V_R_: residual variance.

| **Trait** | **^1^h^2^** | **pe^2^** | **ind^2^** | **^1^V_A_** | **V_ID_** | **V_YR_** | **V_R_** |
| --- | --- | --- | --- | --- | --- | --- | --- |
| Chest girth | 0.30 (0.07-0.65) | 0.28 (0.08-0.69) | 0.59 (0.28-0.89) | 0.001 (<0.001-0.002) |  | 0.001 (<0.001-0.006) | 0.002 (<0.001-0.003) |
| Hind leg length | 0.28 (0.07-0.54) | 0.44 (0.17-0.83) | 0.72 (0.45-0.93) | <0.001 (<0.001-0.001) |  | 0.001 (<0.001-0.005) | 0.001 (<0.001-0.001) |
| Ingest-free body-fat | 0.17 (0.01-0.81) | 0.06 (0.01-0.27) | 0.23 (0.03-0.87) | 0.014 (0.001-0.067) |  | 0.005 (<0.001-0.027) | 0.061 (0.011-0.093) |
| Weight | 0.23 (0.03-0.72) | 0.20 (0.04-0.59) | 0.43 (0.13-0.89) | 0.003 (<0.001-0.001) |  | 0.003 (<0.001-0.013) | 0.007 (0.001-0.011) |
| Start day | ^a^0.07 (0.01-0.21) ^b^0.04 (0.01-0.10) | 0.87 (0.65-0.98) | 0.94 (0.80-0.99) | ^a^0.001 (<0.001-0.002) ^b^13.37 (3.83-34.53) | 0.001 (<0.001-0.002) | 0.019 (0.003-0.081) | 0.001 (<0.001-0.002) |
| End day | ^a^0.10 (0.01-0.31) ^b^0.05 (0.01-0.14) | 0.79 (0.50-0.97) | 0.90 (0.70-0.99) | ^a^0.001 (<0.001-0.003) ^b^17.49 (4.53-46.58) | 0.001 (<0.001-0.003) | 0.011 (0.002-0.048) | 0.001 (<0.001-0.002) |
| Distance | 0.19 (0.01-0.73) | 0.47 (0.03-0.87) | 0.66 (0.17-0.91) | 0.023 (0.001-0.095) | 0.051 (0.001-0.115) | 0.006 (<0.001-0.033) | 0.040 (0.011-0.102) |
| Duration | ^a^0.30 (<0.01-0.93) ^b^0.10 (<0.01-0.31) | 0.45 (0.02-0.98) | 0.75 (0.08-1.00) | ^a^0.083 (0.001-0.302) ^b^1.48 (0.05-19.44) | 0.104 (0.001-0.315) | 0.024 (0.001-0.132) | 0.067 (0.001-0.265) |
| Movement rate | 0.76 (0.24-0.94) | 0.12 (0.01-0.58) | 0.89 (0.65-0.97) | 0.287 (0.092-0.443) | 0.016 (0.001-0.206) | 0.016 (0.001-0.085) | 0.042 (0.012-0.130) |

^1^For Poisson distributed traits (start day, end day, duration) heritability and additive genetic effect values are given in both the latent model state ^a^ and true data state ^b^
